# Identification of Fibrinogen as a Plasma Protein Binding Partner for Lecanemab Biosimilar IgG: Implications for Alzheimer’s Disease Therapy

**DOI:** 10.1101/2024.05.01.591892

**Authors:** Jean-Pierre Bellier, Andrea M. Román Viera, Caitlyn Christiano, Juliana A. U. Anzai, Stephanie Moreno, Emily C. Campbell, Lucas Godwin, Amy Li, Alan Y. Chen, Sarah Alam, Adriana Saba, Han bin Yoo, Hyun-Sik Yang, Jasmeer P. Chhatwal, Dennis J. Selkoe, Lei Liu

## Abstract

**Objective:** Recombinant monoclonal therapeutic antibodies like lecanemab, which target amyloid beta in Alzheimer’s disease, offer a promising approach for modifying the disease progression. Due to its relatively short half-life, Lecanemab, administered as a bi-monthly infusion (typically 10mg/kg) has a relatively brief half-life. Interaction with abundant plasma proteins binder in the bloodstream can affect pharmacokinetics of drugs, including their half-life. In this study we investigated potential plasma protein binding interaction to lecanemab using lecanemab biosimilar.

**Methods:** Lecanemab biosimilar used in this study was based on publicly available sequences. ELISA and Western blotting were used to assess lecanemab biosimilar immunoreactivity in the fractions human plasma sample obtained through size exclusion chromatography. The binding of lecanemab biosimilar to candidate binders was confirmed by Western blotting, ELISA, and surface plasmon resonance analysis.

**Results:** Using a combination of equilibrium dialysis, ELISA, and Western blotting in human plasma, we first describe the presence of likely plasma protein binding partner to lecanemab biosimilar, and then identify fibrinogen as one of them. Utilizing surface plasmon resonance, we confirmed that lecanemab biosimilar does bind to fibrinogen, although with lower affinity than to monomeric amyloid beta.

**Interpretation:** In the context of lecanemab therapy, these results imply that fibrinogen levels could impact the levels of free antibodies in the bloodstream and that fibrinogen might serve as a reservoir for lecanemab. More broadly, these results indicate that plasma protein binding may be an important consideration when clinically utilizing therapeutic antibodies in neurodegenerative disease.

## Introduction

The recent successful trials of monoclonal therapeutic antibodies against amyloid-beta (Aβ) not only slowed symptoms but appeared to alter the biological course of Alzheimer’s disease (AD), lowering pathological forms of Aβ, tau protein, and glial fibrillary acidic protein ^1,2^. Lecanemab, which received full FDA approval in July 2023, is a humanized IgG1 monoclonal antibody raised against synthetic Aβ protofibrils bearing the Arctic APP missense mutation (APP E693G)^3^. Data from a large Phase 3 clinical trial (CLARITY AD) showed reduced amyloid burden by PET scan and slower cognitive decline in patients treated intravenously with 10 mg per kilogram of body weight every 2 weeks ^2^

The binding of therapeutic large and small molecules to proteins in the plasma (plasma protein binding, PPB) can have a profound effect on the pharmacokinetics of therapeutic drugs. After a drug is administered, a portion may bind to proteins (or lipids) while some remains free in the plasma ^4^. Albumin and alpha-acid glycoprotein are common PPBs known to significantly affect the clinical efficiency of small drug molecules ^5^. PPB may help explain the bioavailability of a drug at its target site in tissue, as free drug concentration in blood may reflect free drug concentration in tissue and the pool of drug available to cross the blood-brain barrier ^6^. At the same time, PPB may also serve as a “reservoir” by protecting labile drugs, which might otherwise be quickly degraded or excreted. For example, this phenomenon has been observed for specific classes of antibiotic drugs, significantly extending their half-life, which, in turn, permits a reduction in their dosage and thereby enhances their clinical efficacy ^4^.

While commonly studied for small molecule therapeutics, PPB to therapeutic monoclonal antibodies has been relatively understudied. At present, very little information on PPB for lecanemab is known, and we address this gap in the present report. Using Lec-bs IgG, which should be essentially identical to Lecanemab, and employing equilibrium dialysis, our observations suggested the existence of PPBs for Lec-bs IgG. We then identified fibrinogen as a likely protein responsible for Lec-bs IgG PPB through a combination of size exclusion chromatography, Western blotting, and ELISAs. Together, these results support the significance of fibrinogen as a PPB partner for lecanemab and highlight how PPB may be an important consideration for the use of lecanemab and perhaps other therapeutic antibodies.

## Materials and Methods

### Participants

Data for this study were derived from human plasma samples from 15 participants in the Davis Memory and Aging Cohort (MAC) at Brigham and Women’s Hospital. The Davis MAC is a cross-sectional study recruiting adults 45 years or older, who are either cognitively unimpaired, have mild cognitive impairment, or have mild dementia. The 15 participants were randomly chosen, and a summary of their demographics is presented in Table 1. Among participants, one was diagnosed with Alzheimer’s disease, another one with Parkinson’s disease; the 13 other participants were not diagnostic with neurological diseases. All participants provided informed consent prior to the completion of any study procedures, and all study procedures were reviewed by the Institutional Review Board for MassGeneralBrigham.

**Table 1:** Population demographic.

| **Gender**<br>**(frequency)** |  | **Age**<br>**(year)** |  | **MMSE**<br>**(score)** |  |
| --- | --- | --- | --- | --- | --- |
| Female | 12 | Mean | 63.47 | Mean | 27.5 |
| Male | 3 | SD | 12.14 | SD | 2.43 |

### Materials

Lec-bs IgG was purchased from ProteoGenix (Schiltigheim, France). A list of the immunoreagents used in this study is presented in Table 2. PBS was from Gibco (Grand Island, NY). For the biotinylation of Lec-bs IgG, the Pierce Antibody biotinylation Kit for IP (Thermo Fisher Scientific, Waltham, MA) was used according to the manufacturer’s instructions. Briefly, 100 μg of IgG was reacted with NHS-PEG_4_-biotin in 40-fold molar excess for 30 minutes at RT, then desalted using Zeba 7 kDa spin columns (Thermo Fisher Scientific). The protein concentration of the resulting biotinylated Lec-bs IgG solution was calculated using absorbance at 280 nm and a molar absorption coefficient of 97,860 M^-1^cm^-1^. Fibrinogen (80%) was bought from Millipore Sigma (Burlington, MA) and further purified up to ∼ 95% using size exclusion chromatography (SEC).

**Table 2.** Summary of the immunoreagents used in this study.

| **Antibody/**<br>**reagent** | **Host &**<br>**Clonality** | **Clone ID** | **Company** | **Cat. Number**<br>**(lot)** | **Concentration** |
| --- | --- | --- | --- | --- | --- |
| Lecanemab<br>biosimilar IgG | Human<br>Recombinant<br>monoclonal | NA | Proteogenix<br>(Schiltigheim,<br>France) | PX-TA1746 | ED: 300 µg/ml unlabeled<br>WB: 0.41 µg/ml biotinylated<br>ELISA: 0.82 µg/ml biotinylated |
| Fibrinogen | Mouse<br>Monoclonal | 40F11 | Novus Biological<br>(Centennial, CO) | NB120-10068<br>(22/07-F1-F11) | WB: 0.25 µg/ml |
| Plasminogen | Mouse<br>Monoclonal | 270412 | R&D system<br>(Minneapolis, MN) | MAB2596<br>(UPO032210A) | WB: 1 µg/ml |
| Streptavidin<br>peroxidase |  |  | Jackson<br>ImmunoResearch<br>(West Grove, PA) | 016-030-084<br>(167109) | Dot-Blot: 1 µg/ml |
| Anti-mouse<br>IgG peroxidase | Goat<br>Polyclonal |  | Thermo Fisher<br>(Waltham, MA) | 115-035-146<br>(158821) | WB: 0.1 µg/ml |

### Equilibrium dialysis

The equilibrium dialysis experiments used a 2-chamber Teflon dialysis cell (Harvard Apparatus, Holliston, MA). Each 25 μL compartment was separated with a 300 kDa molecular weight cutoff (MWCO) regenerated cellulose acetate membrane. The experiment started when the “retentate” compartment 1 (C1) was filled with donor human plasma spiked with Lec-bs IgG or phosphate-buffered saline (PBS: 20 mM Sodium Phosphate, 140 mM NaCl, pH 7.4) containing 5 mM EDTA spiked with Lec-bs IgG, and the “filtrate” compartment 2 (C2) was filled with PBS containing 5 mM EDTA. Lec-bs IgG was spiked at 300 mg/ml, a concentration similar to the lecanemab IgG concentration in the blood of lecanemab-treated patients as observed in a pharmacokinetic study of patients receiving multiple doses of the drug ^7^. In some experiments, Lec-bs IgG was spiked into compartment 1 (C1) filled with purified fibrinogen solution (3 mg/ml) in PBS containing 5 mM EDTA, a concentration similar to the average fibrinogen concentration in human plasma ^8^. A protease inhibitor cocktail (cOmplete mini EDTA-free, Roche, Mannheim, Germany) was added to all solutions. Dialysis cells were incubated for 12 hr at room temperature with constant agitation to reach a thermodynamic steady-state, then 1 ml of “retentate” and “filtrate” were dot-blotted onto a nitrocellulose membrane (Bio-Rad Labs, Hercules, CA). Dot blots in which biotinylated Lec-bs was detected were performed as described below, and the signal was quantified using the image densitometry analysis software (GelAnalyzer 23.1). The disequilibrium ratio R_Deq_ were calculated as the modulus of the ratio between the actual value in the filtrate (C2) and its expected ratio at equilibrium (R_Eq_= 0.5), according to the equation: 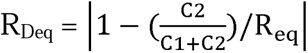.

To test the statistical significance of the disequilibrium, median values for plasma or PBS were compared to the hypothetical median value at equilibrium (R_Deq_= 0) using the one-sample Wilcoxon signed-rank test with a significance level of p= 0.05.

### SEC by fast protein liquid chromatography (FPLC)

SEC was performed by FPLC (AKTApure, Cytiva, Marlborough, MA) using a Superose-6 10-300GL column (Cytiva) to fractionate 0.5 mL of human plasma diluted 1:1 in the running buffer. The running buffer (20 mM Sodium Phosphate, 140 mM NaCl, pH 7.4) flowed at a rate of 0.5 mL/min. Thirty fractions of 0.5 mL each were collected 15 min after injection for subsequent ELISA and Western blot analysis. SEC calibration was performed using a gel filtration standard (Bio-Rad Labs).

### ELISA

For indirect ELISA, SEC fractions were diluted 4-fold in Tris-buffered saline buffer (TBS: 20 mM Tris, 500 mM NaCl, pH 7.55), then 5 μL samples were used to coat an ELISA plate (Nunc, MediSorp, Thermo Fisher Scientific) overnight at 4°C with agitation. Each sample-coated well was incubated with a peroxidase-blocking solution consisting of sodium azide 0.1% and 0.3% hydrogen peroxide in TBS for 1 hr at RT. Then, the wells were washed 3 times and incubated at RT for 1 hr with agitation in a blocking solution of 5% BSA in TBS. After incubation, biotinylated Lec-bs IgG diluted in TBS containing 0.03% Tween-20 (TBS-T) and 1% BSA was added to the wells and incubated at RT for 2 hr with agitation (850 rpm). After incubation, the solution was removed, and the wells were subsequently washed three times before incubation with streptavidin peroxidase at RT for 1 hr with agitation. After 3 washes with TBS-T, 50 μL of 1-step Ultra TMB-ELISA (Thermo Fisher Scientific) was added and incubated for 10 min, then 10% sulfuric acid was added to stop the reaction, and optical density (OD) was measured at 450 nm. For sandwich ELISA, unlabeled Lec-bs IgG was used as the capture antibody, and anti-fibrinogen antibody was used as the detector (Table 2). Fifty microliters of unlabeled Lec-bs IgG diluted in TBS was used to coat an ELISA plate (Nunc, MediSorp, Thermo Fisher Scientific) overnight at 4°C with agitation. The wells were then incubated at RT for 1 hr in a blocking solution of 5% BSA in TBS with agitation. SEC fractions were diluted 4 times in TBS buffer, and 50 μL was added to the wells for 2 hr at RT. The detector antibody (anti-human fibrinogen) was added for 1 hr at RT. A secondary labeled antibody against mouse IgG was added for 1 hr at RT. Fifty microliters of 1-step Ultra TMB-ELISA was added and incubated for 10 min, then 10% sulfuric acid was added to stop the reaction, and OD was measured at 450 nm. A wash step with TBS-T was performed between all steps using a BioTek 405TS auto-washer (Agilent technologies, Santa Clara, CA). A two-way ANOVA analysis combined with the Tukey test was used for statistical analysis.

### Polyacrylamide gel electrophoresis (SDS-PAGE) and Western Blotting (WB)

Reductive and non-reductive SDS-PAGE were used to analyze the SEC fractions from plasma donors. Twenty microliters of SEC fraction was diluted in loading buffer (NuPAGE LDS Sample buffer, Invitrogen), denatured for 5 min at 95°C and electrophoresed on a 4-12% Bis-Tris Sure PAGE (GenScript, Piscataway, NJ) with the molecular weight marker (Prime-Step Prestained, broad range protein Ladder, BioLegend, San Diego, CA) in MES-SDS running buffer (Genscript) at 125V for 45 min. Electrophoresed proteins were transferred at 11V for 70 min to a nitrocellulose membrane (Bio-Rad Labs) using a semi-dry electroblotting method, as previously reported ^9^. Membranes were blocked using 10% skim milk (RPI Corp, Mount Prospect, IL) in 50 mM Tris buffer at pH 7.4 containing 0.15 M NaCl and 0.03% Tween-20 for 1 hr at RT. After blocking, biotinylated Lec-bs IgG was detected by incubating the membranes in streptavidin-peroxidase (1:1000 in same diluent, Jackson ImmunoResearch Labs, West Grove, PA). The membrane was developed using ECL Western blotting substrate (Thermo Fisher Scientific) before digitized analysis of the signal using the Chemidoc Imaging system (Bio-Rad Labs).

### Surface plasmon resonance (SPR)

The equilibrium dissociation constants (K_D_) of Lec-bs IgG for Aβ peptide and fibrinogen were evaluated by SPR using an Affinité Instruments P4SPR (Montréal, Canada). Lec-bs IgG (20 mg/ml in 10 mM acetate buffer, pH 4.5) was immobilized on an Au+ AffiCoat sensor (Affinité Instruments) freshly functionalized using EDC: NHS solution (400 mM: 100 mM in H_2_O).

Binding assays were performed after removing unbound IgG and blocking the reactive site on the sensor. Different concentrations of Aβ peptide of residues 2-42 prepared in 100 mM PBS as previously described ^10^ or purified fibrinogen from human blood prepared in 100 mM PBS + 0.5% BSA were injected, and data were collected.

## Results

### Lec-bs IgG binding to components of human plasma

To investigate whether Lec-bs IgG binds to proteins from the blood, we conducted equilibrium dialysis experiments using human plasma samples from 15 donors. The principle of the equilibrium dialysis experiment is briefly explained in Figures 1A and 1B. In control experiments (as described in Fig. 1A), C1 was filled with PBS spiked with Lec-bs IgG, and C2 was filled with PBS only. After 12 hr of incubation at RT, no statistically significant disequilibrium was measured between compartments (Fig. 1C, “PBS” column, p = 0.125). This experiment showed that a 300 kDa MWCO semi-permeable membrane was suitable for free diffusion of the Lec-bs IgG between compartments and that the incubation time was sufficient to reach a thermodynamic steady state. When compartment C1 was filled with donors’ plasma spiked with biotinylated Lec-bs IgG at 300 mg/ml and then incubated for 12 hr, the absolute values of the disequilibrium ratio (R_Deq_) of the samples ranged from 1.55 % to 40.47 % (Fig. 1C, “plasma” column). A statistically significant disequilibrium ratio (p < 0.0001) between compartments C1 and C2 was observed, suggesting that Lec-bs IgG bound to plasma components.

**Figure 1.**
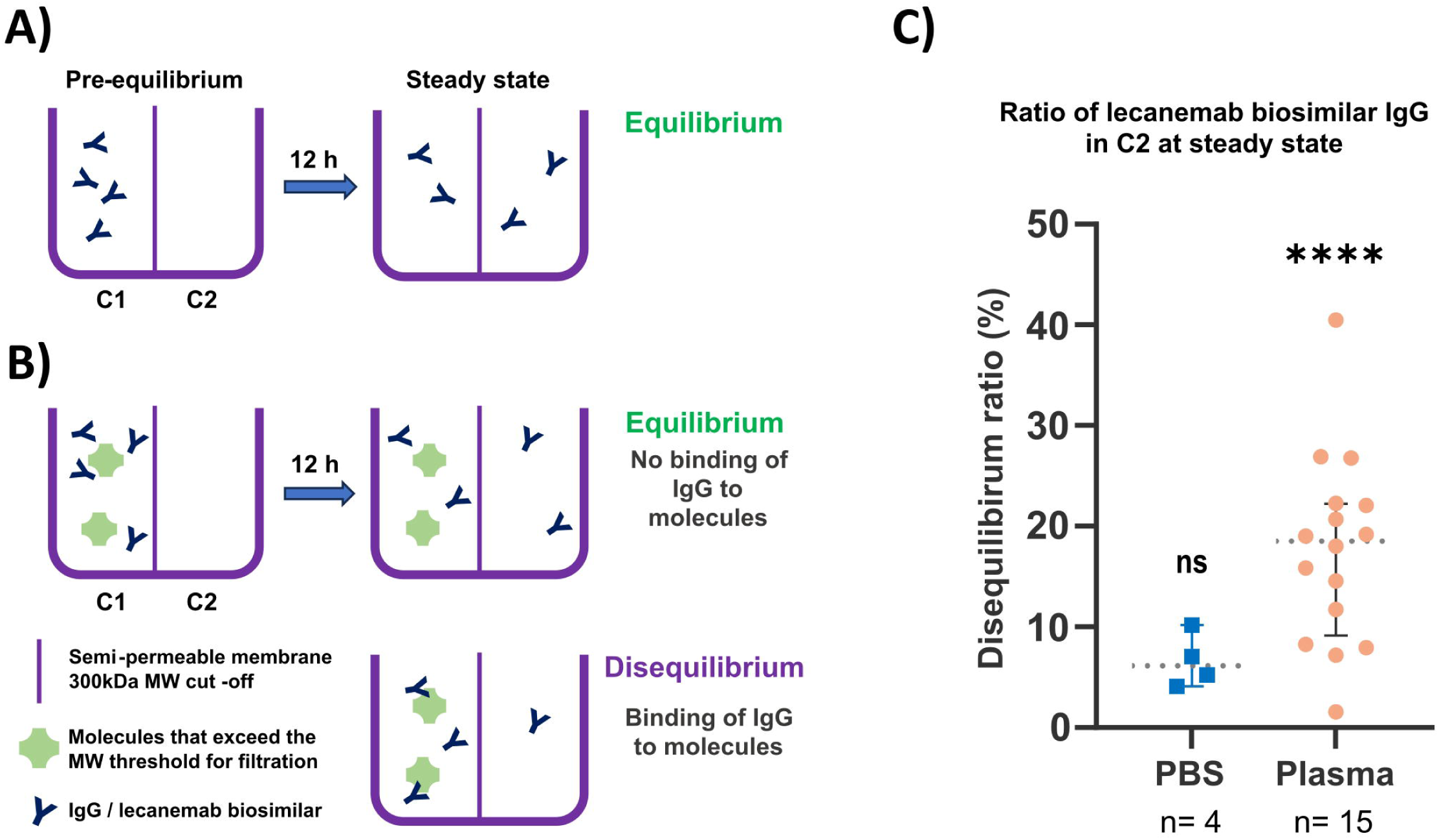
**A)** Equilibrium dialysis experiments comprise a semipermeable membrane with an MWCO of ∼300 kDa that separates two liquid-proof compartments of equal size. In a simple experimental set up with only Lec-bs IgGs, the experiment started with a pre-equilibrium state when compartment 1 (C1) was filled with PBS containing IgG, and compartment 2 (C2) was filled with PBS only. Driven by Brownian motion, IgGs will freely diffuse into C2 because their MW (150 kDa) is lower than the MWCO of the semipermeable membrane. After several hours of incubation at RT, a thermodynamic steady state is reached, and the same concentration of IgG is observed in both compartments. **B)** In another experimental setting, Lec-bs IgGs are in solution with another large molecule (green-colored) that cannot diffuse freely into C2 due to a molecular size exceeding the MWCO of the semipermeable membrane. Schematics on the right show the possible steady states reached after several hours of incubation at RT. When IgGs do not bind to macromolecules, IgG concentrations in C1 and C2 are equal, and equilibrium is observed. When IgGs bind to the macromolecule, IgG concentration in C1 is higher than in C2, and a disequilibrium is observed. **C)** Graph showing disequilibrium ratios of biotinylated Lec-bs IgG between C1 and C2. “PBS” and “Plasma” columns show ratios after 12 hr incubation when IgGs are spiked in PBS (n=4) or in human donor’s plasmas (n=15), respectively. Error bars indicate the 95% CI, and dotted lines indicate the median value. ns: non-significantly different from the ratio at equilibrium (0%). (****: p < 0.0001).

### Lec-bs IgG binding to a high molecular weight (MW) species in human plasma

To further characterize potential Lec-bs IgG binding partners in human plasma, donors’ plasmas were fractionated by SEC using a high-resolution Superose-6 column. The performance of such columns allows the fractionation of large proteins and complexes. Calibration of the columns with globular protein standards showed a wide fractionation range for molecules from 1.3 kDa to 670 kDa (Supp. Fig. 1). The elution profile of the human plasma recorded by absorbance at 280 nm (OD_280nm_) resolved several peaks (Fig. 2B). When aliquots of human plasma SEC fractions were coated onto 96-wells plate and analyzed by ELISA using Lec-bs IgG as a detector antibody, significant immunoreactive peaks were observed (Fig. 2C), with the most robust immunoreactivity detected in high MW fractions (> 670kDa). The strongest immunoreactive peak eluted in fractions 14-15 (black dot in Fig. 2C) and had a MW estimated at 1 MDa by extrapolation from the calibration standards (insert in Fig. 2A). This peak matched an OD_280nm_ peak (black dot in Fig. 2B), indicating a minor but relatively abundant species. Other weaker immunoreactive peaks were observed in fractions of low MW. These observations indicated several plasma proteins bound to Lec-bs IgG but, preferentially, a plasma protein with a native MW > 670 kDa.

**Figure 2.**
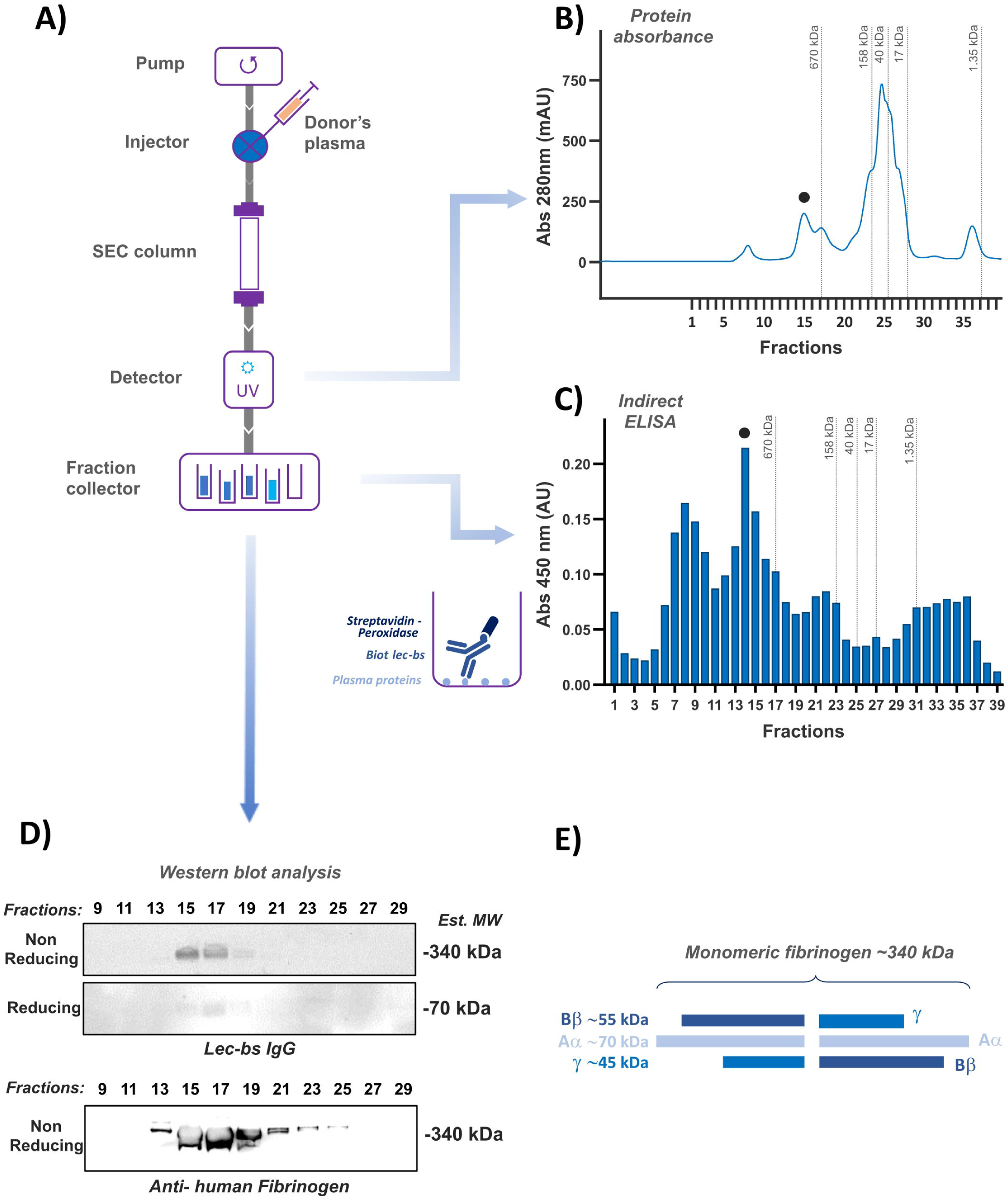
Identification of Lec-bs IgG-immunoreactive human donor’s plasma fractions after size exclusion chromatography (SEC: Superose-6 10-300GL column). **A)** Schematic of the SEC system equipped with a UV detector to monitor UV-absorbance, and a fraction collector. **B)** SEC profile of a typical donor’s plasma as recorded by the UV detector. The collected fractions are indicated as upward ticks on the x-axis. The black dot indicates the position of the highest Lec-bs IgG-immunoreactive peak seen in C). **C)** Indirect ELISA using Lec-bs IgG on the human plasma SEC fractions; the black dot indicates the position of the most immunoreactive peak. **D)** Western blot using Lec-bs IgG (upper boxes) and anti-human fibrinogen (lower box) after electrophoresis of every other SEC fraction (from 9 to 29) by SDS-PAGE in non-reducing or reducing conditions. **E)** Schematic diagram of fibrinogen structure.

### Lec-bs IgG binding to a multimeric protein complex in human plasma

The most robust Lec-bs IgG immunoreactivity fractions showed a high MW, suggesting it could represent a protein complex. To test this hypothesis, SEC fractions 9-29 were analyzed using non-reducing and reducing SDS-PAGE followed by WB for Lec-bs IgG (Fig. 2D). Under non-reducing conditions, immunoreactive bands exceeded the highest MW of the gel’s protein ladder markers (>240 kDa). However, their size was estimated to be ∼340 kDa by extrapolation from the migration pattern of the molecular weight markers. Under reducing conditions, a weak immunoreactive band was observed at a MW of 70 kDa; this discrepancy in signal intensity between non-reducing and reducing conditions suggests that Lec-bs IgG may recognize a conformational epitope. In the present experimental context, the conformational epitope recognized by Lec-bs might require interaction between multiple complexes, suggesting that Lec-bs may preferentially bind a multimeric protein. The strongest immunoreactivity was observed in SEC fractions 15 and 17 in both non-reducing or reducing conditions. Weaker signals were also observed in adjacent fractions. These results fit the strongest immunoreactive peak in fraction 14-15 observed by ELISA (Fig. 2C, black dot). As for other SEC fractions immunoreactive by ELISA, no signal was observed on WB, probably because these fractions contained proteins with MW that are out of the range of the protein size (10-400 kDa) resolved with the SDS-PAGE system used in this study (4-12% Bis-Tris gel + MES-SDS buffer).

### Fibrinogen as a potential binding candidate to Lec-bs IgG

Among the known high-abundance plasma proteins, the above observations fit well with the characteristics of soluble fibrinogen, the third most abundant protein found in human plasma and a known PPB for other drugs. Fibrinogen is a large ∼340 kDa hexameric glycoprotein made of 3 pairs of subunits assembled by disulfide bonding (Aα: 66 kDa, Bβ: 52 kDa, and γ: 46 kDa), which together define a fibrinogen monomer (Fig. 2E). Under reducing conditions, WB using Lec-bs IgG revealed an immunoreactive protein with an MW of ∼70 kDa protein, compatible with the reported 66 kDa of the Aα subunit of fibrinogen. Under non-reducing conditions, WB using Lec-bs IgG revealed a 340 kDa immunoreactive protein similar to the reported MW of fibrinogen. Under native conditions (non-denaturing and non-reducing), as encountered when performing ELISA on SEC fractions, Lec-bs IgG immunoreactivity was observed in fractions eluting from 670 kDa to about 1 MDa, which is compatible with the size of fibrin homodimers (680 kDa) or fibrin degradation products from clot (up to 960 kDa), the most common soluble forms of fibrinogen/fibrin in human plasma ^11^. These observations suggested fibrinogen as a potential PPB to Lec-bs IgG. To verify this hypothesis, a WB using an anti-human fibrinogen antibody was performed on the same SEC fractions of plasma as used previously. The anti-human fibrinogen antibody used was the mouse monoclonal antibody clone 40F11, which recognizes soluble plasma fibrinogen ^12^. Western blot analysis with this anti-human fibrinogen antibody (Fig. 2D) showed immunoreactive bands that display similar patterns and sizes to those observed with WB using Lec-bs IgG.

### Binding of Lec-bs IgG to fibrinogen

To further test the hypothesis that fibrinogen is a potential binding partner to Lec-bs IgG, purified plasma fibrinogen (mostly soluble plasma dimeric fibrinogen) from human blood was subjected to SDS-PAGE under non-reducing conditions. Using purified human fibrinogen, a WB using Lec-bs IgG revealed a single band with an MW estimated at ∼340 kDa (Fig. 3A, middle lane). Using human plasma, a WB using Lec-bs IgG revealed a band with a slightly lower MW (Fig. 3A, right lane). Purified human plasminogen was used as a control and showed no signal with Lec-bs IgG (Fig. 3A, left lane). WB using an anti-human fibrinogen antibody showed identical results (Supp. Fig. 2). In addition, we performed a sandwich ELISA using Lec-bs IgG as a capture antibody and anti-human fibrinogen antibody as the detector antibody coupled with an anti-mouse IgG peroxidase-labeled antibody (see inset in Fig. 3B). Using purified fibrinogen as an analyte, we observed a robust, highly significant immunoreactive signal compared to the control experiments where the analyte was plasminogen (Fig. 3B). For further confirmation, an equilibrium dialysis experiment was performed in which a solution of purified human fibrinogen was spiked with biotinylated Lec-bs IgG. We observed a statistically significant disequilibrium ratio, indicating that Lec-bs IgG interacts with purified human fibrinogen (Fig. 3C). These observations confirmed that Lec-bs IgG was binding to fibrinogen.

**Figure 3.**
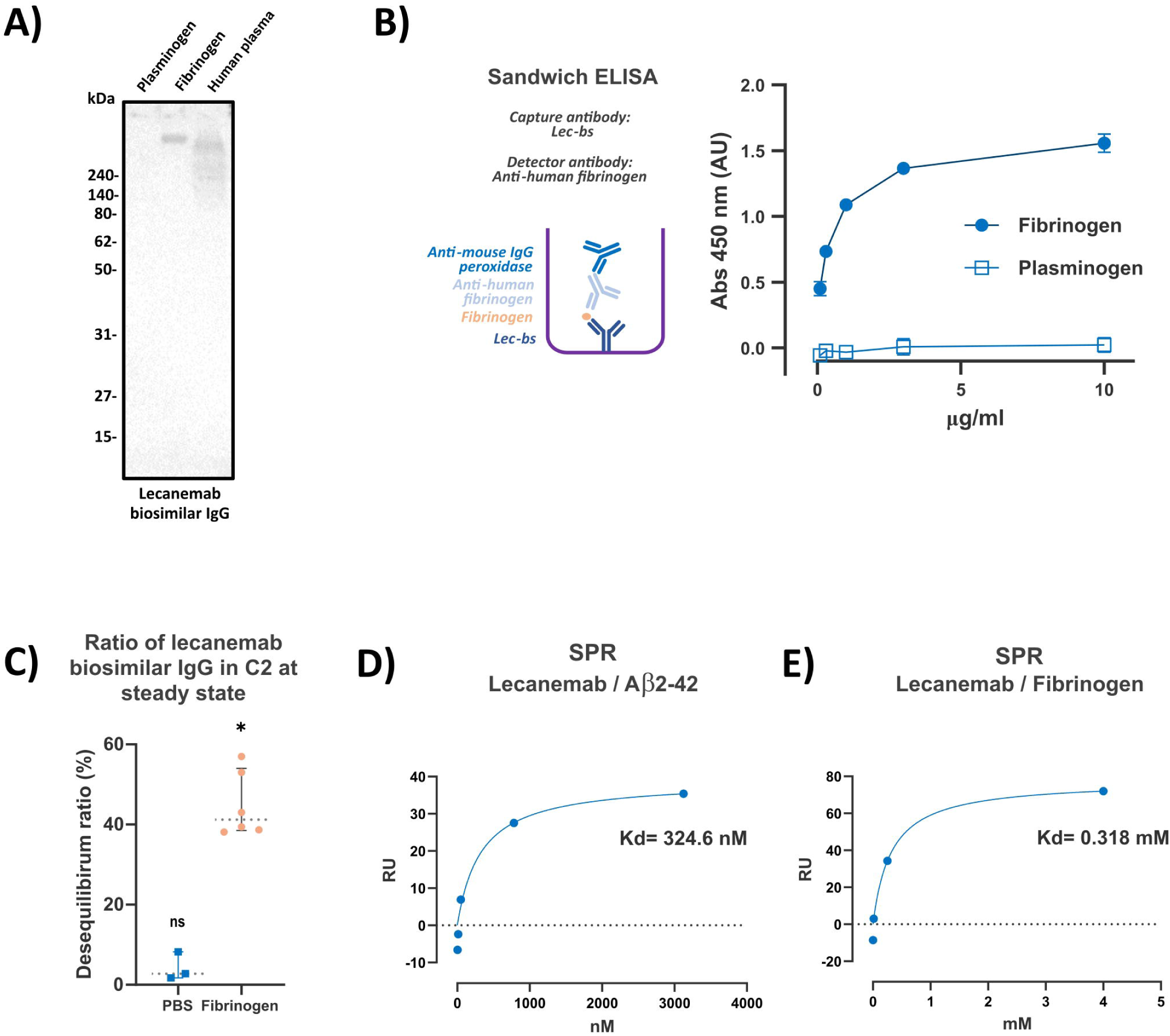
Binding of Lec-bs IgG to purified fibrinogen. **A)** WB using Lec-bs IgG on samples of purified plasminogen, purified fibrinogen, and donor’s plasma. Lec-bs IgG immunoreactive bands are observed only in lanes containing purified fibrinogen and donor plasma. **B)** Sandwich ELISA for purified human fibrinogen and plasminogen using lecanemab as a capture antibody and anti-human fibrinogen antibody as the detection antibody coupled with an anti-mouse IgG peroxidase-labeled antibody. A dynamic response is observed with purified fibrinogen, which is statistically significant and different from the control (purified plasminogen) at any concentration (#: p < 0.0001). **C)** Graph showing disequilibrium ratios of biotinylated Lec-bs IgG between C1 and C2 after 12 hr of incubation (as in Fig. 1) when IgGs were spiked into PBS (n= 3) or in a purified fibrinogen solution (n= 6). Error bars, 95% CI; dotted lines, median value. Non-significantly different from equilibrium (ns); significantly different from equilibrium (*, p = 0.0312). **D)** Determination of K_D_ of lecanemab for Aβ2-42 peptide using SPR. **E)** Determination of K_D_ of lecanemab for human fibrinogen using SPR.

### Low-affinity binding of Lec-bs IgG to fibrinogen

The binding properties of Lec-bs IgG to Aβ and fibrinogen were investigated using SPR. Lec-bs IgG binding properties for Aβ were assessed using a synthetic Aβ peptide of amino acids 2 to 42. This peptide covers almost all the Aβ42 sequence but is less susceptible to spontaneous aggregation. Injection of Aβ monomers over the immobilized Lec-bs showed that Aβ 2-42 bound Lec-bs IgG with a K_D_ of 0.32 μM. These results were similar to those previously reported (i.e., 2.3 μM) for lecanemab with another Aβ species (Aβ 1-28) ^13^. Next, the binding properties of Lec-bs IgG for fibrinogen were assessed by injecting a solution of purified human fibrinogen over immobilized Lec-bs IgG. The apparent affinity of lecanemab for fibrinogen was estimated at 318 mM. These observations indicated that the binding of Lec-bs IgG to fibrinogen is substantially weaker than that to Aβ monomers.

## Discussion

In the present report, we used a biosimilar of the clinical lecanemab to address the possibility that the therapeutic anti-Aβ monoclonal antibody lecanemab demonstrates significant plasma protein binding. Though PPB has been heavily studied in the context of small molecule drugs, the binding of therapeutic antibodies to plasma proteins has been understudied. Using a combination of equilibrium dialysis, SEC, and SDS-PAGE/WB, we observed that Lec-bs showed significant binding to a high abundance plasma protein that we then identified as a multimeric assembly of fibrinogen. Using SPR, we demonstrate that Lec-bs binds to purified fibrinogen with moderate affinity -- approximately 1,000 times less than its affinity for Aβ2-42 monomers. Given the high abundance of fibrinogen and the potential for variability in fibrinogen levels over time and across individuals, these results may be relevant to the clinical use of lecanemab.

Small molecules are commonly tested for PPB phenomena using a “shift” assay in which plasma is spiked with a small drug into an equilibrium dialysis device using a semi-permeable membrane with relatively low MWCO (<30 kDa), allowing free diffusion of the small molecule (e.g., Smith et al., 2010). The size of a therapeutic antibody (∼150 kDa) requires an adaptation of this approach, and we used a semi-permeable membrane with a MWCO of 300 kDa, which allows free diffusion of IgG across the two compartments of the equilibrium dialysis device. Other proteins with a MW lower than the MWCO of the semi-permeable membrane might also diffuse. Using such adaptation of the “shift” assay, we have successfully observed a difference in Lec-bs IgG concentration in normal human plasma between compartments of the equilibrium dialysis device, demonstrating that Lec-bs IgG binds to plasma macromolecular components. Disequilibrium ratios show a relatively wide distribution, suggesting multiple PPBs to lecanemab might exist simultaneously in plasma. To our knowledge, this work provides a rare example of IgG binding to blood components. Antibodies are usually freely soluble in blood; there is no report of binding to serum albumin and few to other components. In some cases of autoimmune disease, IgGs are directed against specific blood components ^14,15^ or when a recombinant antibody was engineered or selected to bind specifically to a blood component to increase its half-life ^16^.

We discovered that fibrinogen, a highly abundant plasma protein, can serve as PPB partner for Lec-bs IgG. Plasma has a highly complex mixture of proteins reflecting polypeptides synthesized in the whole body and relevant to individual physiological and pathophysiological states ^17^. When characterizing possible PPB using ELISA on SEC fractions of donor’s plasmas, we observed the strongest immunoreactivity in a fraction with MW > 670 kDa correlating with a relatively abundant plasma protein. The human plasma proteome comprises thousands of proteins, and this count is still expanding; however, only 20 proteins account for 97% of human plasma protein ^17,18^. The most abundant proteins in plasma include albumin, globulins (α, β, γ), and fibrinogen. Among these highly abundant proteins, the number of relatively abundant proteins with an MW greater than 670 kDa is very limited ^17,19^. Assembled multimeric fibrinogen is among these very few high MW and high abundance proteins, which helped prompt our close examination of fibrinogen and as a PPB partner for Lec-bs. In humans, fibrinogen is the most abundant coagulation factor, accounting for about 7% of the total plasma proteins. Fibrinogen is a large 340 kDa hexameric glycoprotein made of 3-pairs of dimeric subunits assembled by disulfide bonding (Aα: 66k Da, Bβ: 52 kDa, and γ 46 kDa), which together define a monomer. Fibrinogen is synthesized in the liver and it circulates in the blood as a soluble homodimer (680 kDa) ^20^. Upon the action of thrombin, the removal of small fragments (fibrinopeptides) converts fibrinogen into fibrin. Fibrin monomers quickly polymerize into soluble protofibrils, which then aggregate into fibers (the fibrin-clot) essential for hemostasis. Fibrin clots are degraded by plasmin into fibrin degradation products (up to 960 kDa), which are also soluble ^11,21^. Fibrinogen homodimers and fibrin degradation products have size characteristics that fit the size characteristics of the relatively highly abundant protein species that displayed immunoreactivity to Lec-bs IgG in SEC fractions. Therefore, we hypothesized that fibrin(ogen) (soluble homodimer fibrinogen species and small fibrin degradation products) in the donors’ plasmas was a possible Lec-bs IgG PPB candidate. We used WB, ELISA, and SPR analysis to show that Lec-bs IgG binds to purified human fibrinogen (mostly soluble homodimer fibrinogen). These observations demonstrated that fibrinogen is a PPB for Lec-bs IgG. Of note, at least one prior report suggests that fibrinogen may bind to IgG ^22^, though we are not aware of recent examples where such binding has been tested. Given the experimental context, the identification of fibrinogen as a PPB for Lec-bs IgG has been facilitated by its relatively high abundance and structural characteristics (high MW multimeric protein complex) that provide valuable insights. However, our experiments did not rule out the potential existence of other PPBs. For instance, we did not eliminate the possibility of small fibrin degradation products as binding partners for Lec-bs IgG.

We identified fibrinogen based on its molecular characteristics and subsequently confirmed its binding to Lec-bs IgG by sandwich ELISA, WB, and SPR. Most importantly, we estimated the binding affinity value of Lec-bs IgG to fibrinogen by SPR as 0.318 mM. This affinity is three orders of magnitude lower than the binding affinity of Lec-bs IgG to Aβ monomer (K_d_= 0.324 μM). Low binding affinity is a common feature among PPBs. The K_D_ of PPB for small molecules can be as low as 7.7 mM for albumin binding to ceflacor ^23^ or as high as 0.162 nM for albumin binding to bilirubin ^24^, and an average binding affinity could be estimated to approximatively 500 mM. Lec-bs IgG interaction with fibrinogen falls into this average; for example, it has a binding affinity similar to the drug ceftriaxone to human serum albumin (K_d_= 0.316 mM) ^23^. Common PPBs for small-molecule drugs include albumin, α-acid glycoprotein, and lipoproteins. Fibrinogen is a less common PPB, but its ability to contribute to disequilibrium in “shift” assays that test small-molecule drugs has been well documented ^25–27^. These findings are consistent with the fact that fibrinogen exhibits many binding sites for small molecules, including heparin, flavonoids, benzothiazole, and various antibiotics ^26,28–31^, and for proteins such as thrombin, complement receptor 3, plasmin, and Aβ ^21,32,33^. Numerous studies have shown that biosimilars, including monoclonal therapeutic antibody biosimilars, demonstrate high similarity with their reference biological drug ^34–36^. There is no experimental evidence that the Lec-bs IgG is biologically and therapeutically equivalent to genuine, clinical lecanemab IgG. However, we showed that the Kd of Lec-bs IgG to monomeric Aβ (K_d_= 0.324 μM) is in a similar order to what was previously reported for genuine lecanemab IgG ^13^. Therefore, it is reasonable to suggest from our collective data that fibrinogen is a PPB to (genuine) lecanemab.

The efficacy of a drug is related to the free (unbound) drug concentration rather than the total (bound and unbound) concentration of the drug circulating in the patient’s bloodstream ^5^. When highly abundant in blood, a PPB can effectively reduce free drug availability and can have a major impact on the PK/PD of a drug ^37^. In humans, fibrinogen is the most abundant coagulation factor, accounting for about 7% of total plasma protein content, with an average physiological clinical concentration of 1.5 to 4.5 g/L in plasma ^8^, which is also is 10 million times higher than Aβ concentration in human plasma. In this context, despite the modest binding affinity of lecanemab to fibrinogen, a very large amount of fibrinogen in the bloodstream could result in substantial amounts of lecanemab being bound rather than remaining in the free state. In turn, the reduction in free lecanemab may impact the amount of lecanemab available to cross the blood-brain barrier and have a therapeutic impact on neurons. At the same time, the low binding affinity to fibrinogen suggests that lecanemab might be released from fibrinogen when the concentration of free lecanemab drops in the bloodstream. In this context, fibrinogen might act as a reservoir that may help to maintain a steady concentration of lecanemab in the blood of the patient receiving the drug. As mentioned previously, PPB is a phenomenon that can influence drug availability by altering its half-life in the blood. The half-life of a drug is dependent on its clearance and volume of distribution^6^; binding a drug with a short half-life to a PPB with a long half-life such as albumin might extend the half-life of the drug ^38^. However, in contrast to the relatively long half-life of albumin (21 days; Nilsen et al., 2020), the half-life of fibrinogen is short (4 days), and its degradation pathway is not yet resolved^21,40^.The half-life of circulating immunoglobulin G is roughly 10–21 days^41^. The half-life of both lecanemab and donanemab have been reported to be less than 10 days ^7,42,43^, which is shorter than many other previously studied anti-amyloid antibodies. For example, the reported half-life of aducanumab and gantenerumab are 24.8 and 22 days, respectively^44,45^. The half-life of a therapeutic antibody is often attributed in part to the presence of anti-drug antibodies directed against mouse sequences in the paratope of chimeric monoclonal antibodies ^46^ or the lack of binding to neonatal Fc receptor ^47^. None of these factors have been investigated to understand lecanemab half-life, but our demonstration that lecanemab binds to fibrinogen raises the question of whether this binding might alter lecanemab half-life if lecanemab follows the degradation pathway of fibrinogen. The clinical implications of fibrinogen as a PPB remain unclear, and further work is needed to understand how the interaction of lecanemab with fibrinogen may impact drug dosing and contribute to inter-individual variability in drug response.

## Conclusion

We identify fibrinogen as a major potential plasma protein binder for lecanemab based on its molecular characteristics and relative abundance in plasma. These results support the hypothesis that fibrin(ogen) may be a PPB for lecanemab; however, we cannot exclude the possibility that other plasma proteins may also bind to lecanemab. We found that the K_D_ of Lecanemab to fibrinogen is in the range of the K_D_ observed for known PPB with various drugs. Fibrin(ogen) as a PPB for lecanemab may impact the bioavailability of lecanemab in clinically important ways, including by serving as a reservoir for lecanemab and altering its half-life. Further work is needed to understand these potential clinical implications, including whether fibrin(ogen) function is altered following lecanemab binding and whether plasma fibrin(ogen) levels should be considered in the dosing of lecanemab.

## Acknowledgment

We are grateful to the members of the Selkoe Laboratory for helpful discussion. The authors express gratitude to all blood donors for their generosity. GelAnalyzer 23.1.1 (available at www.gelanalyzer.com) by Istvan Lazar Jr., PhD and Istvan Lazar Sr., PhD, CSc. This work was funded by NIH Grants RF1AG079569 (JPC and LL), R01AG071865 (DJS, JPC, and LL) nd the Davis APP program at Brigham and Women’s Hospital (BWH) (DJS, JPC, and LL). The funders had no role in data collection, analysis, or publication decisions.

## Authors contributions

Conceptualization: JPB, JPC, DJS, LL; Resources: HSK, JPC, DJS, LL; Methodology: JPB, AMR; Blood collection: CC, JAUA, SM, ECC, LG; Investigation: JPB, AMR, AL, AC; Data curation: JPB; Formal analysis: JPB; Supervision: JPB; Funding acquisition: JPC, DJS, LL; Validation: JPB, AMR, SMA, AS, HBY; Visualization: JPB; Project administration: JPB, HSK; Writing – original draft: JPB; Writing – review and editing: JPB, HSK, SMA, JPC, DJS, LL.

