## Supplementary figures for "Identification of Fibrinogen as a Plasma Protein Binding Partner for Lecanemab Biosimilar IgG: Implications for Alzheimer’s Disease Therapy" (from another version of this preprint: version 1, 2024-05-01)

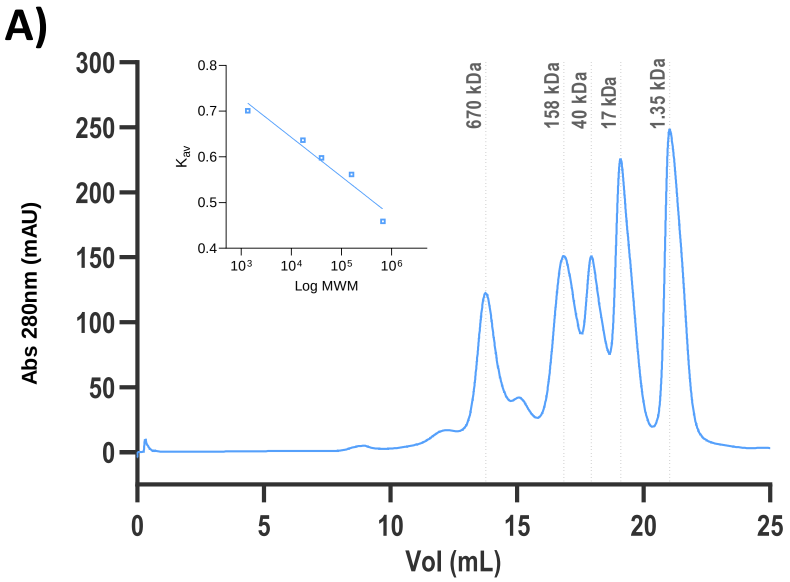


**Supp. 1**

SEC profile and corresponding calibration curve (insert) for the gel filtration protein standards. The MW of the standards is indicated in the graph.

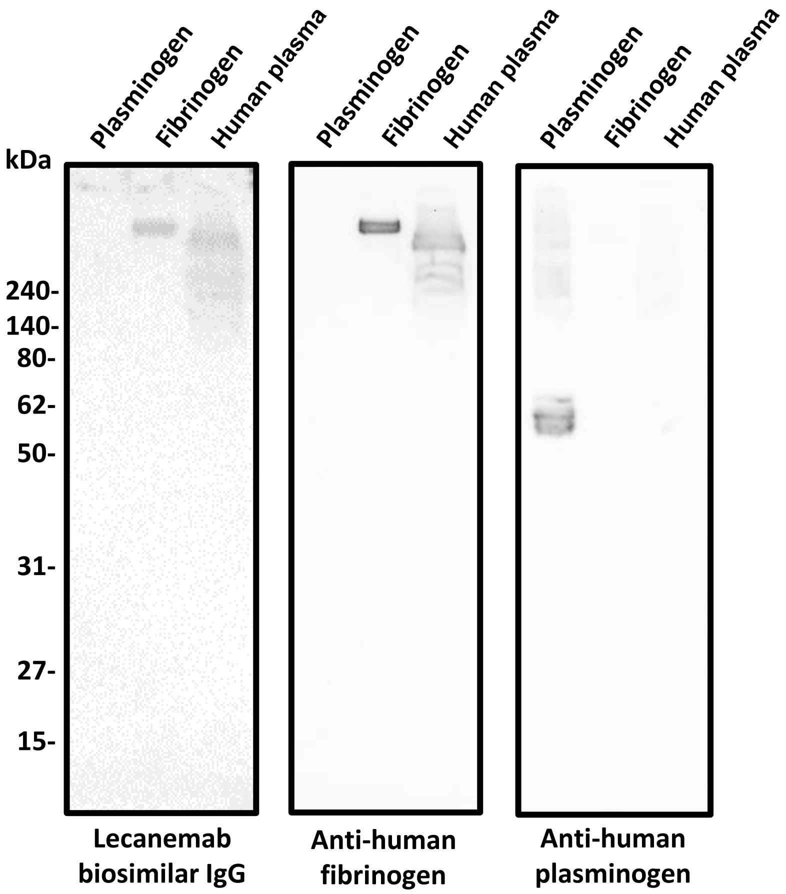


**Supp. 2**

Western blot a sample of purified plasminogen, purified fibrinogen, and donor's plasma using Lecanemab biosimilar IgG (left blot), anti-human fibrinogen (middle blot), and anti-human plasminogen (right blot). The left blot is the same as shown in Figure 3A and shows the binding of lecanemab to purified fibrinogen and patient plasma. Western blot using anti-human fibrinogen confirms that the pattern of lecanemab biosimilar IgG staining is like the pattern of fibrinogen staining (middle blot). Western blot using anti-human plasminogen confirms the identity of the purified plasminogen sample, and the absence of similarity with the pattern of lecanemab biosimilar IgG staining(right blot).
